# Copy-back viral genomes induce a cellular stress response that interferes with viral protein expression without affecting antiviral immunity

**DOI:** 10.1101/2023.05.17.541157

**Authors:** Lavinia J. González Aparicio, Yanling Yang, Matthew S. Hackbart, Carolina B. López

## Abstract

Antiviral responses are often accompanied by translation inhibition and formation of stress granules (SG) in infected cells. However, the triggers for these processes and their role during infection remain subjects of active investigation. Copy-back viral genomes (cbVGs) are the primary inducers of the Mitochondrial Antiviral Signaling (MAVS) pathway and antiviral immunity during Sendai Virus (SeV) and Respiratory Syncytial virus (RSV) infections. The relationship between cbVGs and cellular stress during viral infections is unknown. Here we show that SG form during infections containing high levels of cbVGs, and not during infections with low levels of cbVGs. Moreover, using RNA fluorescent in situ hybridization to differentiate accumulation of standard viral genomes from cbVGs at a single-cell level during infection, we show that SG form exclusively in cells that accumulate high levels of cbVGs. PKR activation is increased during high cbVG infections and, as expected, PKR is necessary to induce virus-induced SG. However, SG form independent of MAVS signaling, demonstrating that cbVGs induce antiviral immunity and SG formation through two independent mechanisms. Furthermore, we show that translation inhibition and SG formation do not affect the overall expression of interferon and interferon stimulated genes during infection, making the stress response dispensable for antiviral immunity. Using live-cell imaging, we show that SG formation is highly dynamic and correlates with a drastic reduction of viral protein expression even in cells infected for several days. Through analysis of active protein translation at a single cell level, we show that infected cells that form SG show inhibition of protein translation. Together, our data reveal a new cbVG-driven mechanism of viral interference where cbVGs induce PKR-mediated translation inhibition and SG formation leading to a reduction in viral protein expression without altering overall antiviral immunity.

**One Sentence Summary:** cbVGs trigger the cellular stress response independent of the antiviral response during RSV and parainfluenza virus infection leading to a reduction of virus protein expression.

## Introduction

Respiratory syncytial virus (RSV) and parainfluenza viruses are endemic RNA viruses responsible for a large disease burden, especially involving children and older adults (1, 2). RNA viruses produce not only full-length standard viral genome (stVG) but also variants, hypermutated RNAs, and non-standard viral genomes (nsVGs) that provide different functions and advantages to the virus (3, 4). nsVGs produced during RSV and parainfluenza virus infections are critical determinants of infection outcome in vitro and in vivo (5–7). When produced early during infection, nsVGs significantly reduce virus spread and disease severity in mice and humans (5, 7). nsVGs impact the infection via stimulation of major signaling pathways that shape the cellular response to the infection. Identifying cellular pathways and molecular mechanisms by which nsVGs reduce virulence may lead to new strategies to prevent severe disease upon RNA virus infection.

One nsVG subpopulation, copy-back viral genomes (cbVGs), has critical roles in inducing the cellular antiviral immune response, controlling the rate of viral replication, and promoting the establishment of persistent infections (6–8). Non-segmented negative-sense RNA viruses generate cbVGs when the viral polymerase initiates replication at the promoter region, falls off the template and then reattaches to the nascent strand (4). The polymerase then uses the nascent strand as a template and continues replicating, copying back the already synthetized RNA (4). The resulting RNA molecules contain highly structured immunostimulatory motifs and lack genes encoding viral proteins (7, 9). Although cbVGs can only replicate in the presence of a full-length standard genome that provides essential viral proteins, cbVGs are key interactors with the host and drive several cellular responses that determine the infection outcome. Notably, all the known effects of cbVGs on shaping the host response are dependent on the Mitochondrial Antiviral Signaling (MAVS) pathway. cbVGs activate Retinoic acid-Inducible Gene I (RIG-I)-like receptors (RLRs) leading to MAVS signaling which then induces robust antiviral responses (9). By activating the MAVS pathway, cbVGs stimulate the interferon response that ultimately reduces virus spread and induces long term protective immunity (6). Additionally, cbVGs signal through MAVS to activate a cell survival mechanism that promotes the establishment of persistent infections in vitro (8). Whether cbVGs can induce other cellular pathways that contribute to the outcome of the infection remains unknown.

In addition to the antiviral immune response, virus infections can induce cellular stress responses that lead to protein translation inhibition and the formation of stress granules (SG) (10). During most viral infections, the cellular stress response is initiated upon activation of the double-stranded RNA binding protein PKR which phosphorylates the eukaryotic initiation factor 2 alpha (eIF2α) leading to cap-dependent translation arrest, disassembly of polysomes and formation of SG. SG are liquid phase-separated non-membranous organelles composed mostly of untranslated mRNA and RNA binding proteins (11, 12). Activation of this cellular stress response during infection can lead to reduced viral protein expression (12) and has been proposed to mediate the antiviral immune response (13–17).

The predicted overlapping antiviral roles of the PKR-driven cellular stress response and cbVGs led us to question if cbVGs are involved in SG formation during RSV and parainfluenza virus infections, and whether cbVG-mediated antiviral immunity depends on SG formation. Our data show that cbVGs are the primary inducers of canonical SG during Sendai virus (SeV) and RSV infections through PKR activation, and that this induction is independent of the MAVS signaling pathway. Contrary to previous reports, we found that MAVS does not localize to cbVG-induced SG and that translation inhibition and SG formation are not required for overall induction of antiviral immunity. Instead, we show that cbVGs induce protein translation inhibition in SG positive cells resulting in reduced levels of virus proteins at a single cell level without affecting the expression of antiviral proteins at a population level. Overall, these data demonstrate that cbVGs orchestrate the induction of cellular stress and antiviral immunity independently, highlighting the importance of considering the presence of nsVGs when studying virus-host interactions. Importantly, our data uncovers a new primary mechanism of interference by cbVGs via the induction of viral protein translational arrest.

## Results

### SG form during RSV infection containing high levels of cbVGs

To assess whether cbVGs induced SG formation, we infected lung epithelial A549 cells with RSV stocks containing high or low levels of cbVGs (RSV cbVG-high and RSV cbVG-low, respectively). To achieve high and low cbVG accumulation in these stocks the virus was grown at different multiplicity of infection (MOI), as virus expansion at high MOI promotes the accumulation of cbVG while virus expansion at low MOI cbVGs reduces the accumulation of cbVGs (18). cbVG contents in the stocks were confirmed by PCR (**Figure S1A**). Because cbVGs potently induce the interferon (IFN) response, we expect cbVG-high stocks to induce higher expression of *IL-29* than a cbVG-low stock (6). As expected, *IL-29* mRNA levels were increased in cells infected with cbVG-high stocks (**Figure S1B**). Additionally, presence of cbVGs during infection is expected to correlate with reduced levels of virus replication in infected cells as compared to cbVG-low stocks due to the activity of IFNs (6). Using *RSV G* mRNA transcripts as a proxy for virus replication, we confirmed that infection with RSV cbVG-high stocks resulted in reduced levels of *RSV G* mRNA as compared to infection with an RSV cbVG-low stocks (**Figure S1B**).

To visualize SG formation during RSV infections, cells were immunostained for the well-characterized SG associated protein Ras GTPase-activating protein-binding protein 1 (G3BP1), along with the RSV nucleoprotein (NP) to identify infected cells. Fluorescence imaging analysis showed SG in infected cells during RSV cbVG-high infections, while they were rarely detected in RSV cbVG-low infections. SG were observed as early as 12 hours post-infection (hpi) and were still present at 24 hpi (**Figure 1A**). The percent of SG positive cells during RSV cbVG-high infection increased over time, and approximately 10% of infected cells were SG positive at 24 hpi (**Figure 1B**).

**Figure 1:**
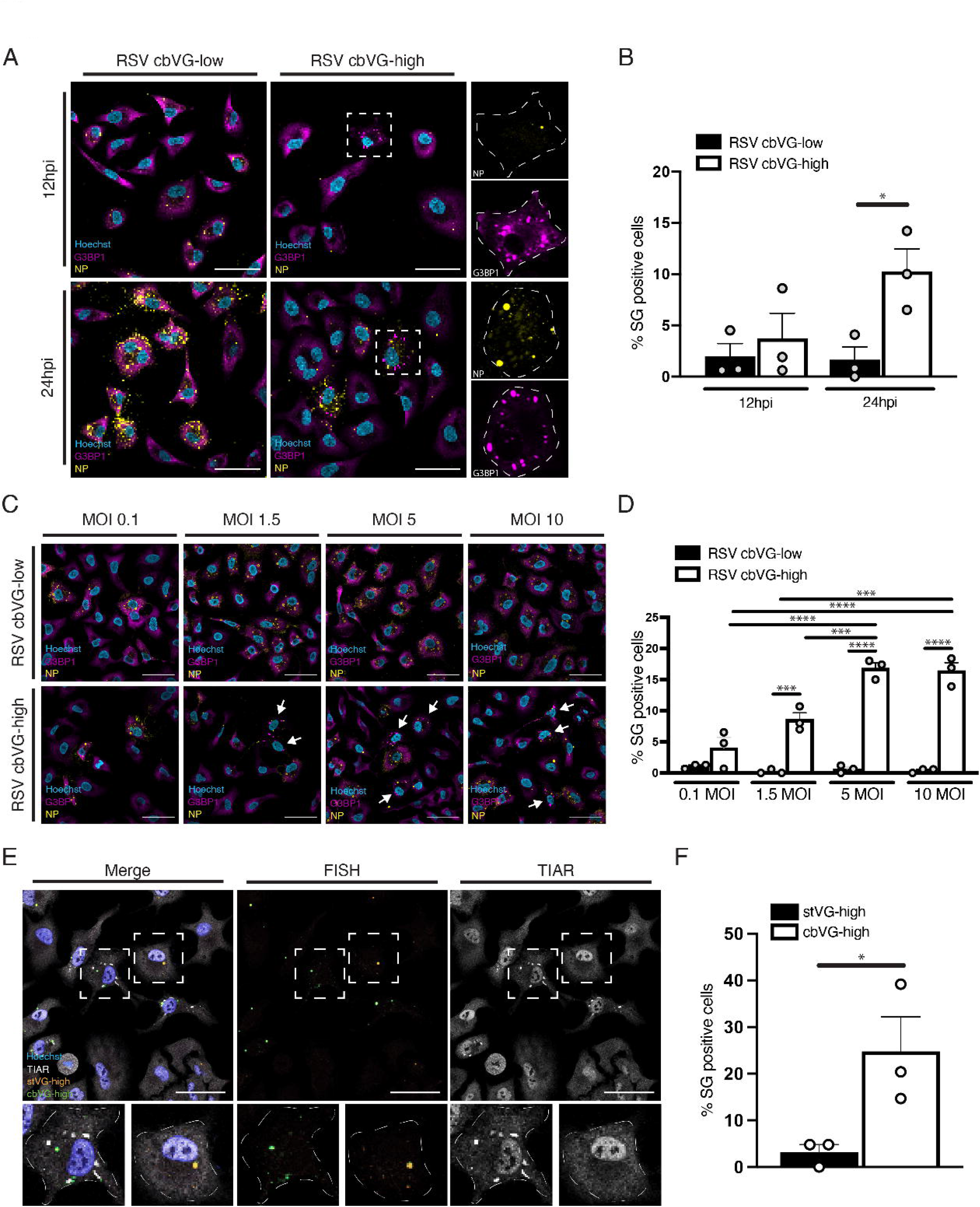
SG form during RSV infection containing high levels of cbVGs. (A) SG (G3BP1, magenta) and viral protein (RSV NP, yellow) detection 12 and 24 hpi with RSV cbVG-high or cbVG-low at MOI of 1.5 TCID_50_/cell. (B) Percent of SG positive cells within the infected population 12 and 24 hpi with RSV cbVG-high and cbVG-low infections. Approximately 150 infected cells were counted per condition (average of three independent experiments shown). (C) SG (G3BP1, magenta) and viral protein (RSV NP, yellow) detection 24hpi with RSV cbVG-high or cbVG-low at MOIs 0.1, 1.5, 5 and 10 TCID_50_/cell. (D) Percent of SG positive cells within the infected population 24 hpi with RSV cbVG-high and cbVG-low infection at MOIs 0.1, 1.5, 5 and 10 TCID_50_/cell. Approximately 150 infected cells were counted per condition (average of three independent experiments shown). (E) SG detection (TIAR, white) in cells staining via FISH for stVG-high (orange) and cbVG (green) cells 24 hpi with RSV cbVG-high at MOI 1.5 TCID_50_/cell. (F) Percent of SG positive cells within the stVG-high and cbVG-high cell populations during RSV cbVG-high infection (average of three independent experiments shown). All widefield images were acquired with the Apotome 2.0 at 63x magnification and are representative of three independent experiments. Scale bar = 50 μm. Statistical analysis: One way ANOVA (*p < 0.05, **p < 0.01, ***p < 0.001, ****p < 0.00001).

Although cbVG-containing viral particles can infect cells, they are not considered fully infectious as they can only replicate in cells co-infected with standard virus particles. Thus, infections based on multiplicity of infection (MOI) only account for the number of fully infectious particles in the inoculum. We expect that RSV cbVG-high infections, which contain both infectious standard particles and non-infectious cbVG particles, will contain a higher amount of total viral particles. To determine if the observed differences in SG formation were due to differences in total viral particles added in the inoculum, we infected cells with RSV cbVG-high and RSV cbVG-low at increasing MOIs and compared percent of SG positive cells. Increasing the MOI of RSV cbVG-low infection did not increase the percent of SG positive cells even when using 10 times more RSV cbVG-low than RSV cbVG-high (**Figures 1C and 1D)**. We observed an increase in the percent of SG positive cells as we increased the MOI during RSV cbVG-high infection, which correlates with the increased number of cbVG-containing particles in the inoculum. However, no differences in percent of SG positive cells were observed between MOI 5 and MOI 10 **(Figure 1D)**, suggesting there is a threshold on the amount of SG positive cells we can obtain at a given time during the infection. Taken together, these data indicate that presence of cbVGs during RSV infection correlates with SG formation.

### SG form exclusively in cbVG-high cells during RSV cbVG-high infection

Using a previously described RNA Fluorescence in situ hybridization (FISH)-based assay that allows differentiation of full-length genomes from cbVGs at a single-cell level (8), our lab reported that cells infected with RSV or SeV cbVG-high stocks have heterogenous accumulation of viral genomes; some cells accumulate high levels of standard genomes (stVG-high) and others accumulate high levels of cbVGs (cbVG-high) (8, 19, 20). To determine if SG formed differentially within these two populations of cells, we combined RNA FISH with immunofluorescence to detect SG during RSV cbVG-high infection. At 24 hpi, SG formed almost exclusively in cbVG-high cells (green) and not stVG-high cells (orange) (**Figure 1E)**. Interestingly only around 30% of the cbVG-high cells had SG (**Figure 1F)**. This could suggest that a threshold of cbVG accumulation in the cells is needed for SG formation or that SG formation occurs asynchronously during infection which is observed during HCV infection (21). Nevertheless, these data demonstrate that cbVGs trigger SG formation.

### cbVGs induce SG during SeV infection

To determine whether cbVG induction of SG also occurs during infection with parainfluenza viruses, we infected cells with cbVG-high or cbVG-low SeV, a member of the paramyxovirus family and close relative to the human parainfluenza virus 1. Like infection with RSV, SG formed predominantly during SeV cbVG-high infections (**Figure 2A)** where approximately 20% of the infected cells were positive for SG at 24 hpi **(Figure 2B).** Compared to cells with undetected SGs or NP (**Figure 2A, right panel inset 1)**, some SG-positive cells had notably low NP signal (**Figure 2A right panel inset 2**) while other SG positive cells showed high NP signal (**Figure 2A, right panel inset 3)**.

**Figure 2:**
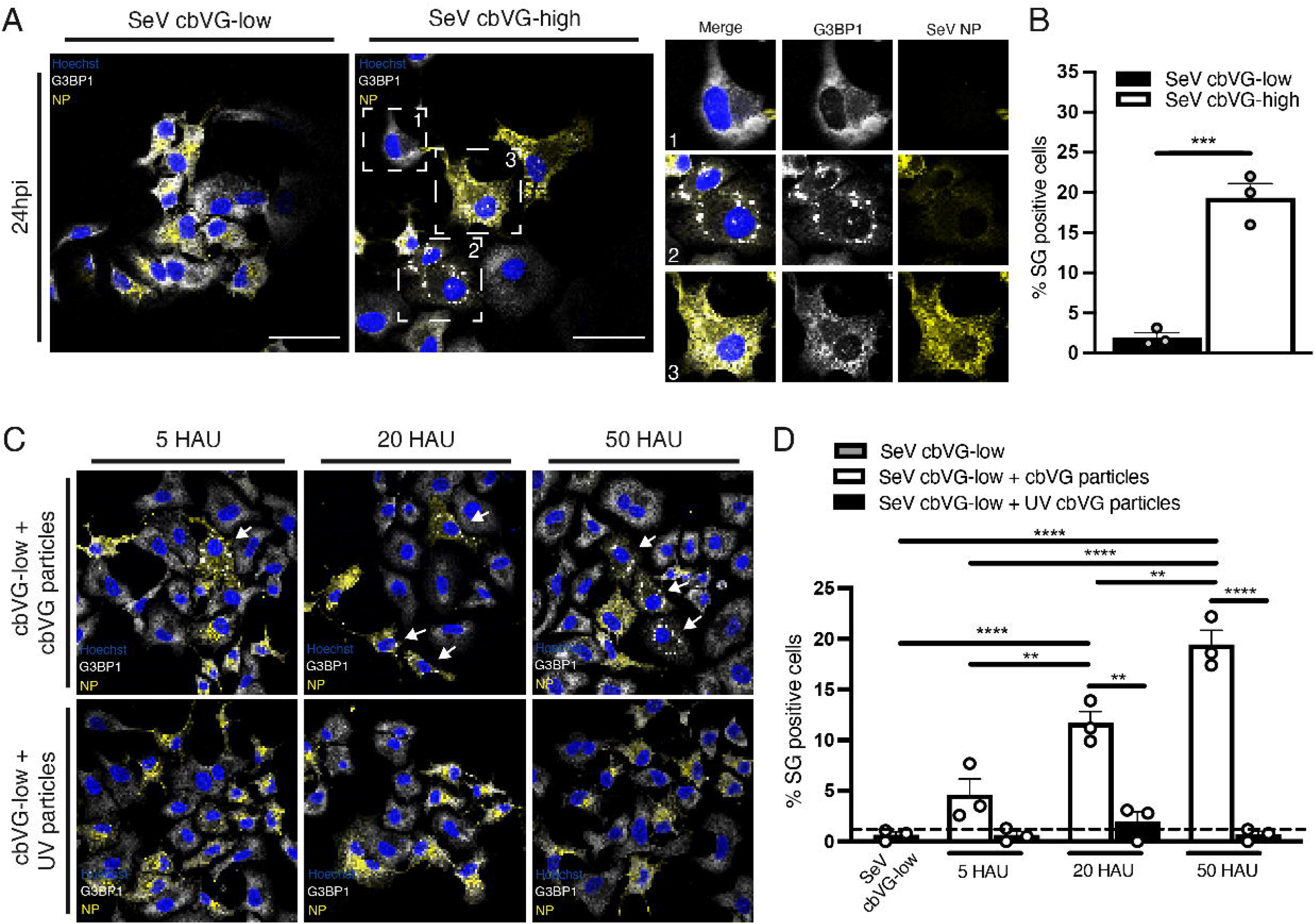
SeV cbVGs induce SG formation. (A) SG (G3BP1, white) and viral protein (SeV NP) detection 24 hpi with SeV cbVG-low and cbVG-high (NP, yellow) at MOI 1.5 TCID_50_/cell. Digital zoomed images for each of the marked cells are shown in the panel on the right. (B) Percent of infected SG positive cells 24 hpi with SeV cbVG-low and cbVG-high at MOI 1.5 TCID_50_/cell. Approximately 150 infected cells were counted per condition (average of three independent experiments shown). (C) SG (G3BP1, white) and viral protein (SeV NP) detection 24 hpi at MOI 1.5 TCID_50_/cell supplemented with either purified cbVG particles or UV-inactivated cbVG particles at increasing hemagglutination units (HAU). (D) Percent of SG positive cells at increasing HAU doses of active/UV inactive cbVG particles. Approximately 200 infected cells were counted per condition (average of three independent experiments shown). All widefield images were acquired with the Apotome 2.0 at 63x magnification and are representative of three independent experiments. Scale bar = 50 μm. Statistical analysis: One way ANOVA (*p < 0.05, **p < 0.01, ***p < 0.001, ****p < 0.00001).

To further establish the role of cbVGs in inducing SG, we performed a dose-dependent experiment using purified cbVG-containing viral particles. We infected cells with SeV cbVG-low and supplemented the infection with increasing doses of purified cbVG-containing particles. The percent of SG positive cells increased in proportion to the amount of purified cbVG-containing particles added **(Figure 2C upper panel and 2D)**. SG were not observed, however, when we added the same amounts of UV-inactivated purified cbVG particles **(Figures 2C lower panel, and 2D)**. These data demonstrate that only replication-competent cbVGs induce SG formation during RNA virus infection.

### Canonical SG are formed during cbVG-high infection

Some viruses can induce formation of SG-like granules that differ compositionally from canonical SG and can re-localize SG components to viral replication centers (22–24). Other viruses induce formation of RNAseL dependent bodies (RLBs), which contain common proteins also found in SG but are structurally and functionally distinct from SG (25). To better characterize the granules observed during RSV cbVG-high infection, we began by testing if cbVG-dependent granules require polysome disassembly, a crucial step for proteins to bind ribosome-free mRNA and form canonical SG. For this we treated RSV cbVG-high infected cells with cycloheximide (CHX) which inhibits canonical SG by preventing polysome disassembly (26). Sodium arsenite, a chemical known to induce canonical SG was used as a positive control (27). Treatment with CHX during RSV cbVG-high infection led to a decrease in SG positive cells compared to treatment with the drug’s vehicle alone (DMSO) (**Figure 3A, 3B**). To rule out any effect the drugs could have on G3BP1 localization, we co-stained with another SG marker, TIA-1 related (TIAR) protein. Co-staining with TIAR showed co-localization with G3BP1 in SG in the DMSO treated cells and disassembly from granules in the drug-treated conditions (**Figure 3A**), demonstrating that cbVG-dependent SG are canonical SG.

**Figure 3:**
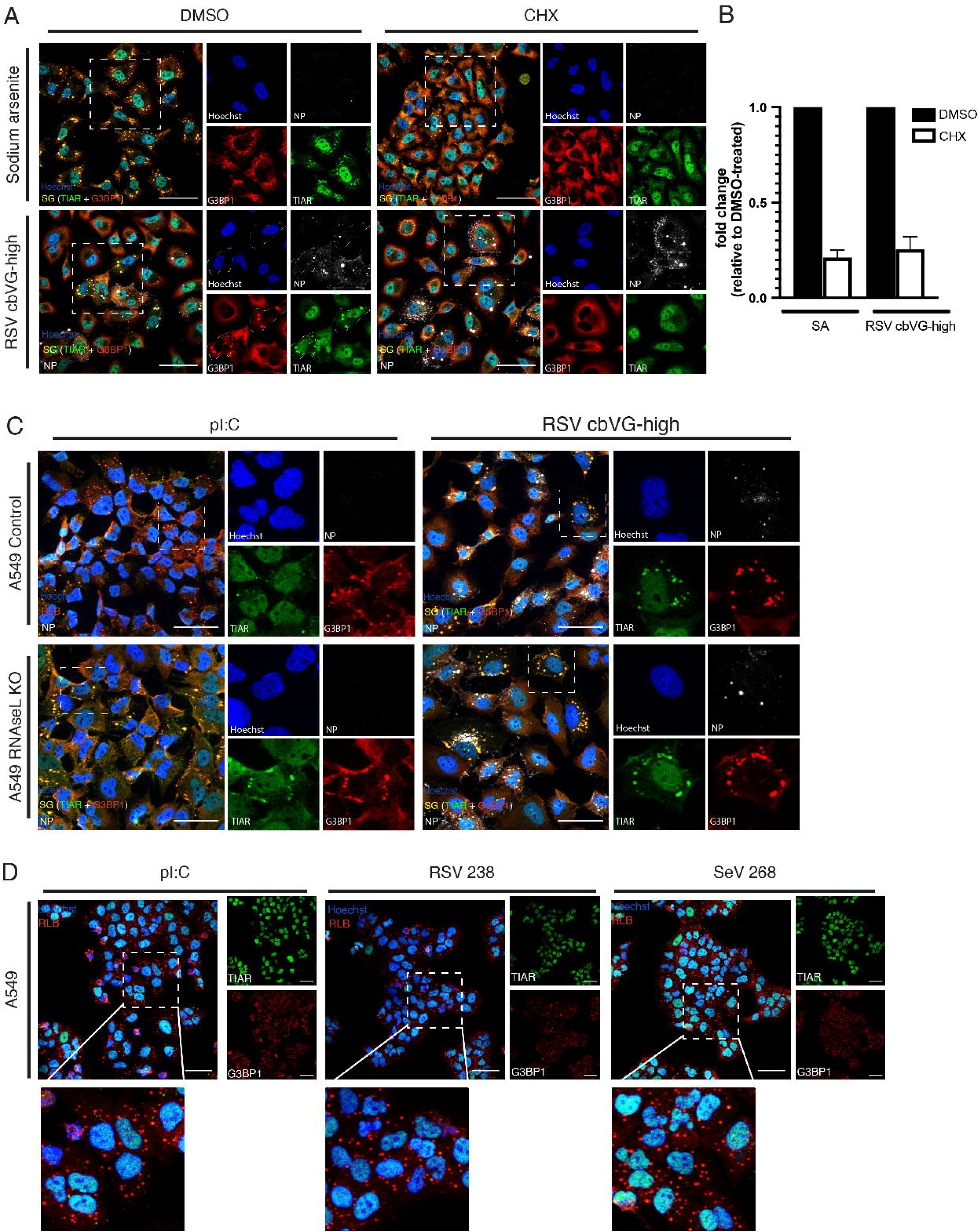
RNA granules formed during RSV cbVG-high infection are canonical SG. (A) G3BP1 (red) and TIAR (green) staining for SG in cells treated with sodium arsenite (SA, 0.5mM) for 1 h or infected with RSV cbVG-high (RSV NP, white) at MOI 1.5 TCID_50_/cell 23 hpi and treated with DMSO, CHX (10 μg/mL), or ISRIB (200nM) for 1 h. (B) Quantification of SG positive cells after drug treatment in sodium arsenite or RSV cbVG-high infected cells. Approximately 150 cells were counted for each condition. Fold change relative to DMSO-treated cells is shown. (C) RNA granule detection (G3BP1, red; TIAR, green) in control and RNAseL KO cells transfected with poly I:C 10 μg/mL or infected with RSV cbVG-high (RSV NP, white) 24hpi at MOI 1.5 TCID_50_/cell. (D) RNA granule detection (G3BP1, red; and TIAR, green) A549 cells transfected with poly I:C or RSV and SeV cbVG derived oligonucleotides RSV 238 and SeV 268. All widefield images were acquired with the Apotome 2.0 at 63x or 40x magnification. Scale bar = 50 μm.

We next tested whether RSV-induced granules were RLBs (28). To do this, we infected RNAseL knockout cells with RSV cbVG-high virus and looked at differences in SG formation comparing to poly I:C transfection which is known to induce RLB formation (28). Structurally, RNAseL dependent bodies (RLBs) are smaller, more punctate, and contain less TIAR than canonical SG (**Figure 3C, left panel)**. RNAseL activation prevents canonical SG from forming by degrading free mRNA necessary for SG to form and only when knocking out RNAseL can canonical SGs form upon stimulation (28, 29). SG are structurally bigger and less uniform than RLBs. SG formed during RSV cbVG-high infection even in RNAseL knockout (KO) cells, and the structure of these granules was unchanged between cell lines, demonstrating that RSV-dependent SG are not RLBs (**Figure 3C**).

We then investigated if, out of the context of an infection, cbVG RNA would still induce formation of canonical SG, or would induce RLBs similar to poly I:C. We transfected in vitro transcribed RSV and SeV cbVG-derived oligonucleotides that maintain the key stimulatory domains of cbVGs (RSV 238 and SeV 268 (9)) into A549 cells and compared to poly I:C-induced RLBs. We saw no differences in RNA granule formation and G3BP1 and TIAR contents between poly I:C RLBs and the granules observed with transfected cbVG-derived oligonucleotides (**Figure 3D**) indicating that cbVG induce canonical SG only in the context of SeV or RSV infection while RLBs are produced in response to naked cbVG RNA.

### cbVG-dependent SG are PKR-dependent and MAVS-independent

To better understand the molecular mechanisms leading to SG formation in response to cbVGs during infection, we investigated the role of major dsRNA sensors in SG induction. SG formation during infection with many viruses, including RSV, depends on PKR activation (11). To confirm cbVGs induce PKR activation, we probed for PKR activation during RSV cbVG-high infection. As expected, PKR is phosphorylated during RSV cbVG-high infections compared to RSV cbVG-low or mock infection **(Figure 4A)**. To determine if cbVG-induced SG are PKR dependent, we infected A549 PKR KO cells (**Figure 4B, middle lane**) and visualized SG formation. Consistent with the literature, PKR KO cells infected with RSV cbVG-high virus did not show SG positive cells (**Figures 4D and E middle panel and bar**). *RSV G* mRNA levels were similar between cell types, confirming that inhibition of SG in PKR KO cells was not due to lower replication of the virus (**Figure 4C, middle bar**). Together, these data suggest that the SG observed during RSV cbVG-high infection are PKR dependent and that cbVG induction of SG is mediated by PKR activation.

**Figure 4:**
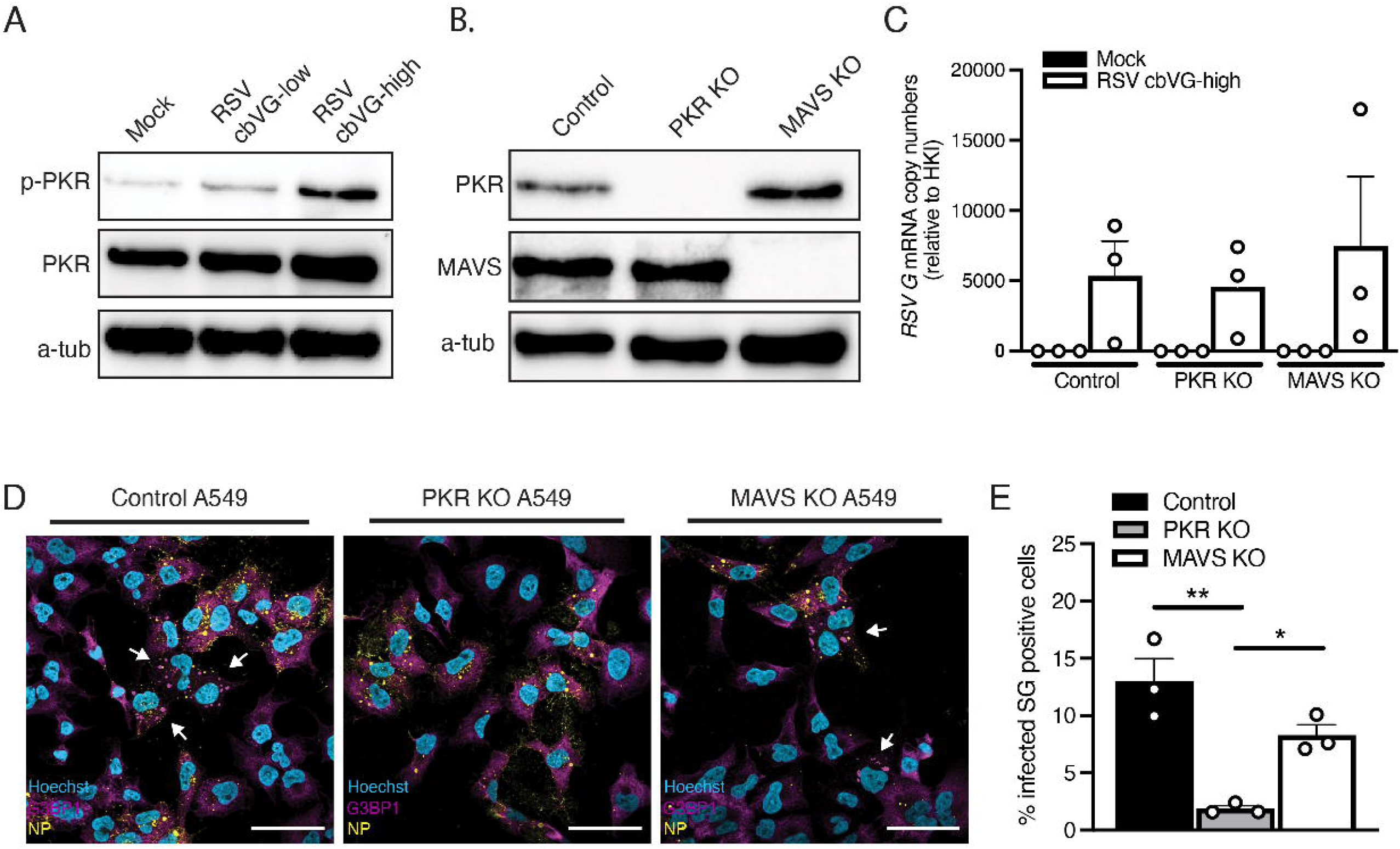
cbVG-dependent SG are PKR-dependent and MAVS-independent. (A) Phosphorylation of PKR 24 hpi with RSV cbVG-low and cbVG-high infection at MOI 1.5 TCID_50_/cell. (B) Western blot analysis showing efficient KO of PKR and MAVS in A549 cells. (C) Expression of *RSV G* gene mRNA relative to the house keeping index (hki) 24 hpi with RSV cbVG-high infection at MOI 1.5 TCID50/cell in control, MAVS or PKR KO A549 cells (average of three independent experiments are shown). (D) SG (G3BP1, magenta) and viral protein (RSV NP) detection in PKR KO and MAVS KO A549 cells 24 hpi with RSV cbVG-high virus at MOI 1.5 TCID_50_/cell. (E) Quantification of SG positive cells 24 hpi with RSV cbVG-high at MOI 1.5 TCID_50_/cell in PKR or MAVS KO A549 cells. Approximately 300 cells were counted per condition. All widefield images were acquired with the Apotome 2.0 at 63x magnification and are representative of three independent experiments. Scale bar = 50 μm. Statistical analysis: One way ANOVA (*p < 0.05, **p < 0.01)

Because cbVGs exert most of their functions through RLR stimulation which leads to MAVS activation and enhanced production of IFN, we sought to investigate whether cbVGs also induced SG through MAVS signaling. To our surprise, MAVS KO cells (**Figure 4B, right lane**) infected with RSV cbVG-high virus showed SG positive cells (**Figure 4D, 4E right panel and bar**). The percent of SG positive cells trended slightly lower than control but was not statistically significant (**Figure 4E).** This is most likely due to a reduced expression of PKR, a well-known interferon stimulated gene (ISG). Contrary to reports in the literature, we did not observe localization of MAVS in SG (**Figure S2A)**, nor recruitment of RIG-I to SG during SeV cbVG-high infection (**Figure S2B).** These data indicate that cbVGs induce SG independent of cbVGs immunostimulatory activity. To our knowledge, this is the first time cbVGs have shown to modulate cellular processes that are independent of MAVS signaling.

### cbVG-dependent SG inhibition is both G3BP1 and G3BP2-dependent

To form SG, nucleating factors initiate RNA protein aggregation and liquid phase-separation (10). Studies suggest that one of these nucleating factors, G3BP1, is necessary and sufficient for SG to form during viral infections (30–32). To determine if G3BP1 is sufficient for cbVG-dependent SG, we infected G3BP1 KO cells (**Figure 5A, second lane**) with RSV cbVG-high virus and looked at SG using TIAR staining as proxy for SG formation. RSV G mRNA levels confirmed that there were not significant differences in viral replication between cell types (**Figure 5B**). Unexpectedly, we observed TIAR-containing SG in G3BP1 KO cells (**Figure 5C, upper panel**). To confirm that these were canonical SG and not aggregation of TIAR as an artifact of knocking out G3BP1, we treated the cells with CHX. Indeed, TIAR-containing SG in G3BP1 KO cells are sensitive to CHX, suggesting these are canonical SG (**Figure 5C, lower panel**). These data indicate that knocking out G3BP1 is not sufficient to inhibit RSV-dependent SG, contradicting what has previously been suggested in the literature (30).

**Figure 5:**
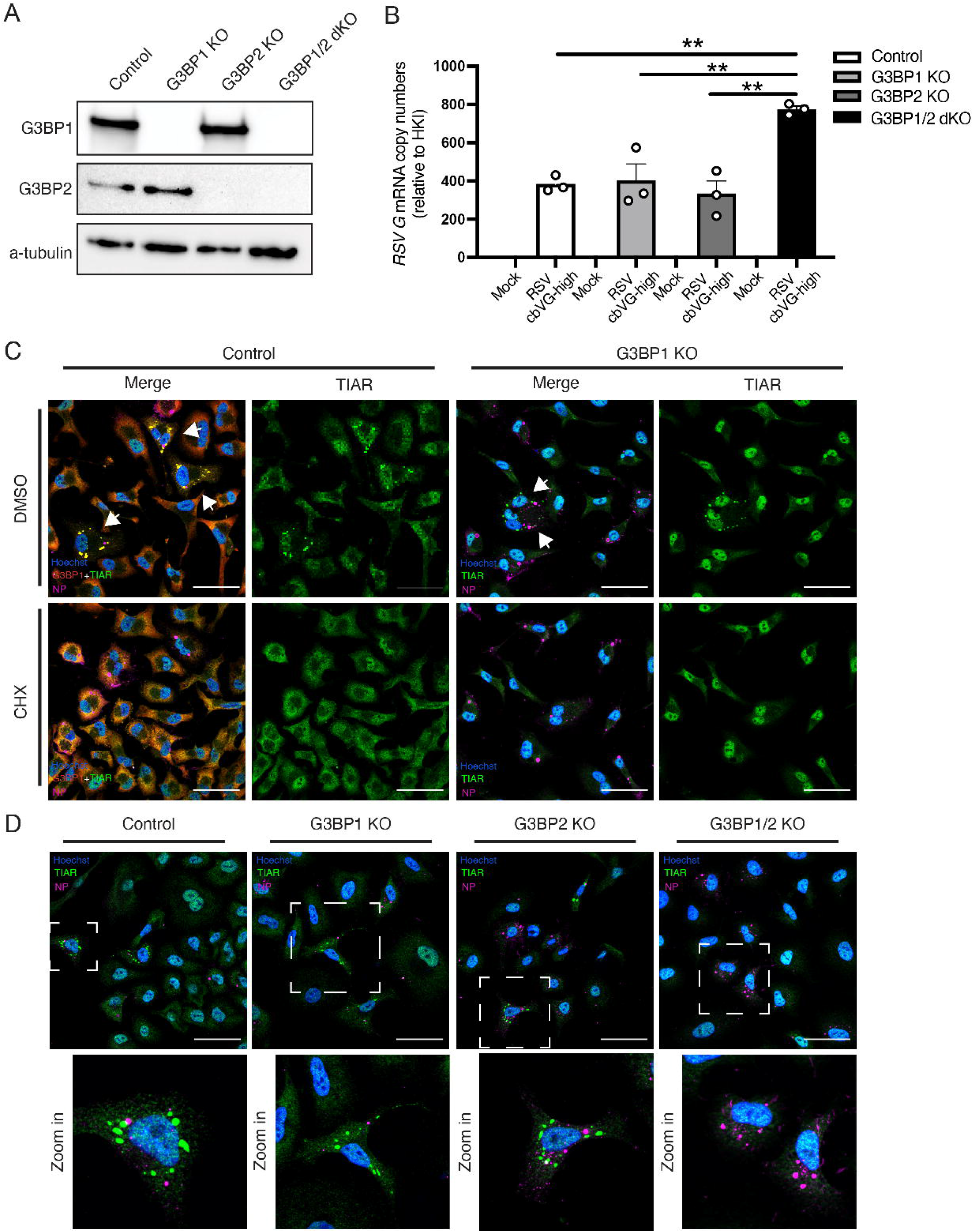
cbVG-dependent SG inhibition is both G3BP1 and G3BP2 dependent. (A) Western blot analysis validating A549 G3BP1 KO, G3BP2 KO and G3BP1/2 dKO (B) Expression of *RSV G* gene mRNA relative to the house keeping index (hki) 24 hpi with RSV cbVG high infection at MOI 1.5 TCID50/cell in control, G3BP1 KO, G3BP2 KO and G3BP1/2 dKO (C) G3BP1 (red) and TIAR (green) staining for SG and viral protein (RSV NP) detection in control and G3BP1 KO cells 24 hpi with RSV cbVG-high at MOI 1.5 TCID_50_/cell and treated with DMSO (upper panel) or CHX (10μg/mL) (lower panel). (D) G3BP1 (red) and TIAR (green) staining for SG and viral protein (RSV NP) detection in control, G3BP1 KO, G3BP2 KO and G3BP1/2 dKO cells infected 24 hpi with RSV cbVG-high at MOI 1.5 TCID_50_/cell. All widefield images were acquired with the Apotome 2.0 at 63x magnification and are representative of three independent experiments. Scale bar = 50 μm. Statistical analysis: One way ANOVA (**p < 0.01).

In the context of some non-virus induced stresses, knocking out both G3BP1 and G3BP2 have shown to be necessary for SG inhibition (33). To test if cbVG-dependent SG inhibition require KO of both G3BP1 and G3BP2, we next generated a G3BP2 KO cell line as well as a G3BP1/2 double KO (dKO) cell line (**Figure 5A**). When we infected G3BP1/2 dKO cells with RSV cbVG-high virus stocks, we no longer observed SG upon staining for TIAR, but SG were still formed in G3BP1 and G3BP2 single KO cells (**Figure 5D**). These data demonstrate that cbVG-dependent SG inhibition requires KO of both G3BP1 and G3BP2.

### cbVG-dependent SG are not required to induce the antiviral response

As SG formation is often associated with induction of the intrinsic antiviral immunity (12, 14, 16, 17), we then determined if SG are necessary for the expression of antiviral genes in response to cbVGs. To do this, we infected control, G3BP1 KO, G3BP2 KO and G3BP1/2 dKO cells with RSV cbVG-high and looked for differences in expression of genes involved in antiviral immunity, including interferons and interferon-stimulated genes (ISGs), at 24 hpi by qPCR. Expression of *IL-29*, *ISG56* and *IRF7* mRNAs was not impaired when comparing control and G3BP1/2 dKO cells, and only statistically significant differences were observed in *IL-29* expression between G3BP1 KO and and dKO (**Figure 6A-C)**.

**Figure 6:**
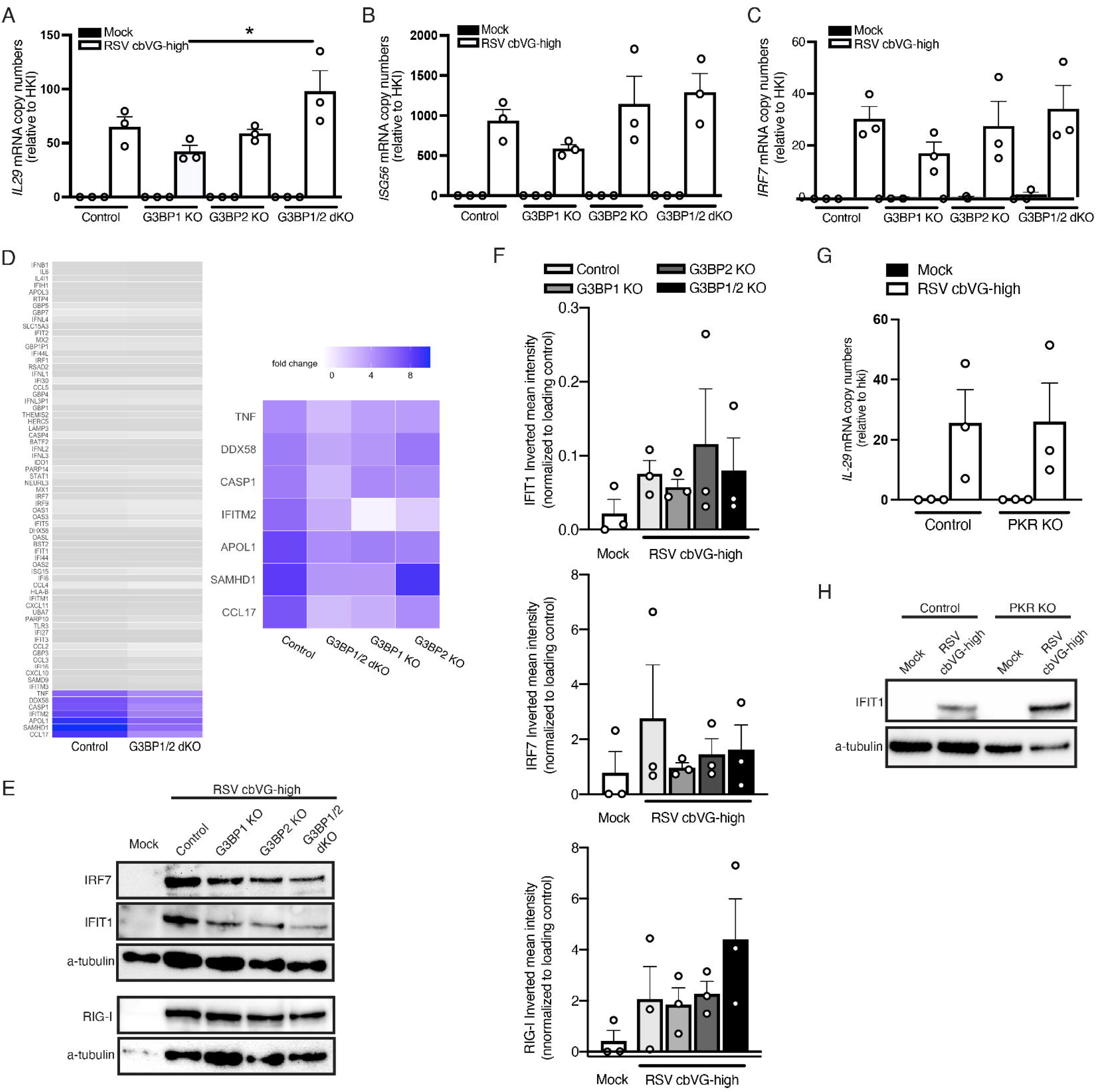
cbVG-dependent SG are not required for the antiviral response during RSV cbVG-high infection. mRNA copy numbers of (A) *IL29*, (B) *ISG56* and (C) *IRF7* relative to the house keeping index (hki) in control, G3BP1 KO, G3BP2 KO and G3BP1/2 dKO cells 24 hpi with RSV cbVG-high at MOI 1.5 TCID_50_/cell. Statistical analysis: One way ANOVA (*p < 0.05) (D) Log 2-fold change analysis of genes related to the antiviral response in control, G3BP1 KO, G3BP2 KO and G3BP1/2 dKO cells 24 hpi with RSV cbVG-high at MOI 1.5 TCID_50_/cell relative to mock infected cells. Genes that had less than a 2-fold decrease difference between control and G3BP1/2 dKO are represented in grey color. Genes that had more than a 2-fold decrease difference are highlighted in color. Genes with 2-fold decrease or more difference between control and G3BP1/2 dKO are shown in the right panel and compared to the log 2-fold change of G3BP1 and G3BP2 single KOs. (E) Western blot analysis of RIG-I, IFIT1 and IRF7 in control, G3BP1 KO, G3BP2 KO and G3BP1/2 dKO cells 24 hpi with RSV cbVG-high at MOI 1.5 TCID_50_/cell. (F) Inverted mean intensity quantification of IRF7, IFIT1 and RIG-I western blot bands relative to α-tubulin loading control. Statistical analysis: One way ANOVA. No statistical significance was found. (G)) *IL29* mRNA levels relative to the house keeping index (hki) in control and PKR KO cells cells 24 hpi with RSV cbVG-high at MOI 1.5 TCID_50_/cell. Statistical analysis: One way ANOVA. No statistical significance was found. (E) Western blot analysis of IFIT1 and IRF7 in control and PKR KO cells 24 hpi with RSV cbVG-high at MOI 1.5 TCID_50_/cell.

To assess the impact of SG on the host antiviral response more broadly, we looked at the whole transcriptome in control and KO cells at 24 hpi. Most ISGs were expressed at similar levels in control and dKO cells (difference in expression were less than 2-fold; **Figure 6D**). In the few cases when there were differences of 2-fold decrease or more in expression, the difference was also observed in the G3BP1 or G3BP2 single KO conditions, suggesting the difference is driven by processes independent of SG formation (**Figure 6D, right panel)**. Additionally, we tested whether absence of SG leads to reduced protein expression of ISGs. Expression of IFIT1, IRF7 and RIG-I was not different between the cell lines, demonstrating that the antiviral immune response is not dependent on SG formation (**Figure 6E, 6F**).

Because the role G3BPs have in the stress response is directly in SG formation and not the translation inhibition that occurs upstream of the pathway, we looked at the direct role of PKR signaling in antiviral immunity. For this, we infected PKR KO cells with RSV cbVG-high virus and compared *IL-29* transcript levels and IFIT1 protein levels to control infected cells and saw no significant differences (**Figure 6G, 6H**). Similarly, cells infected with SeV cbVG-high virus had no differences in phosphorylation of IRF-3, the primary transcription factor leading to type I IFN expression, nor differences in protein expression of the antiviral gene IFIT1 (**Figure S3A, S3B**). Altogether, these data suggest that PKR activation and SG formation are dispensable for global induction of antiviral immunity.

### SeV cbVG-dependent SG form dynamically during infection and correlate with reduced viral protein expression

To study the dynamics of SG assembly and disassembly as well as assess the impact of SG during infection, we generated G3BP1-GFP expressing A549 cells to visualize SG formation in real time. Using live-cell imaging of cells infected with a recombinant SeV expressing miRFP670 and supplemented with purified cbVG particles, we show dynamic formation and disassembly of SG throughout the course of the infection (**Video 1**). During the period of 6 – 72 hpi we identified several subpopulations of cells (**Figure 7A**). Some cells formed SGs after infection and eventually disassembled them (**Figure 7A, series 1).** These cells showed faint levels of miRFP670 signal early in infection. Once SG disassembled, the miRFP670 signal increased. Other cells formed SG and eventually died (**Figure 7A, series 2**). A few cells assembled and disassembled SG and remained very low in miRFP670 signal throughout the infection (**Figure 7A, series 3**). Moreover, formation of SG persisted in the population even 13 dpi **(Figure 7B).** These data demonstrate that SeV cbVG-dependent SG form asynchronously and that formation of SG continues throughout the infection.

**Figure 7:**
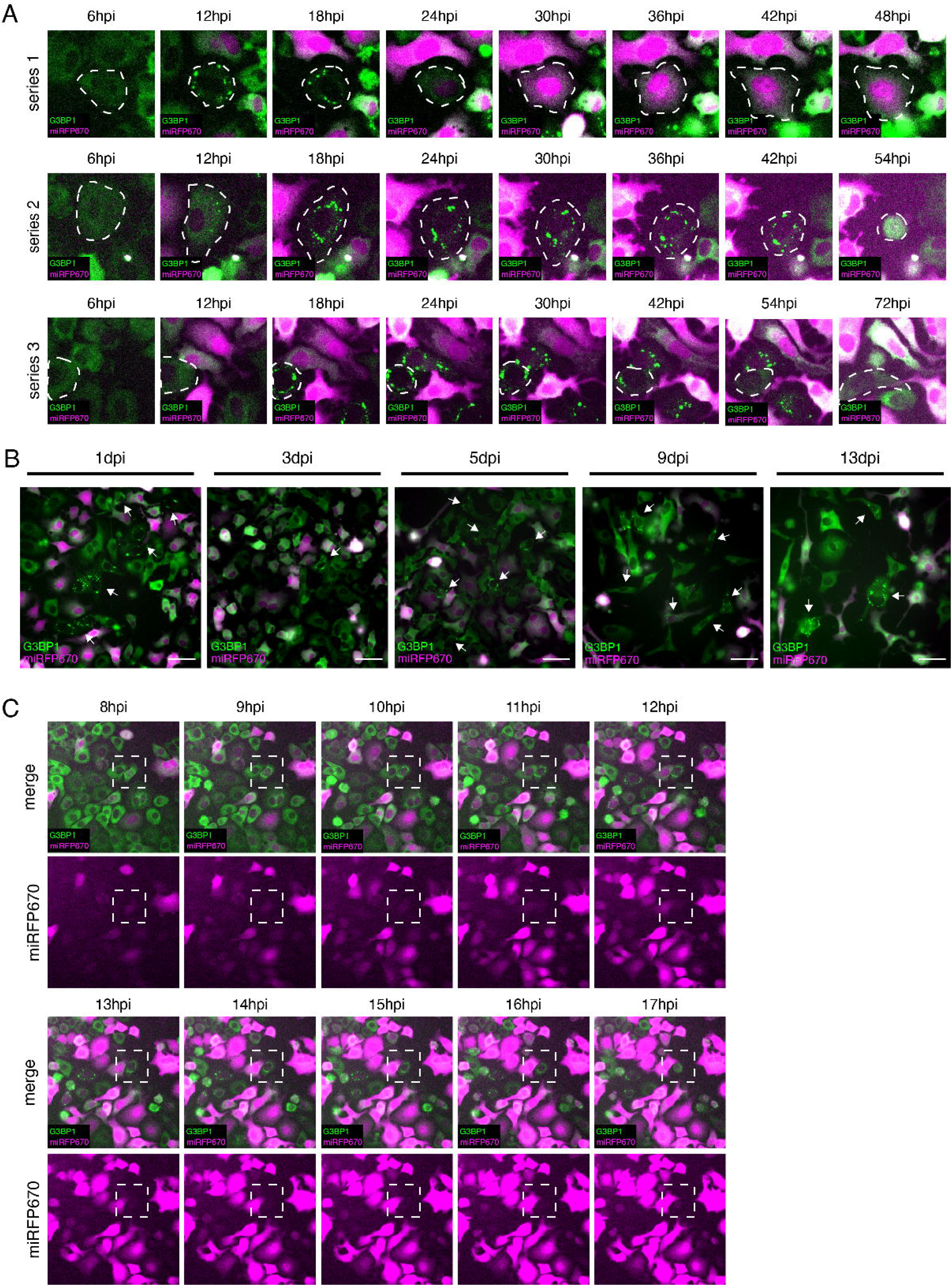
SeV cbVG-dependent SG form asynchronously and are maintained at the population level throughout the infection. (A) G3BP1-GFP (green) expressing A549 cells infected with rCantell-miRF670 (magenta) reporter virus at MOI 3 TCID_50_/cell with 20 HAU of supplemented cbVG purified particles, timelapse microscopy 6 - 72hpi, images every 6 h at a 20x magnification. Series show focus of different cells in the population (B) timelapse microscopy images of G3BP1-GFP (green) expressing A549 cells infected with rCantell-miRF670 (magenta) reporter virus at MOI 3 TCID_50_/cell with 20 HAU of supplemented cbVG purified particles from day 1 to day 13 (C) G3BP1-GFP (green) expressing A549 cells infected with rCantell-miRF670 (magenta) reporter virus at MOI 3 TCID_50_/cell with 20 HAU of supplemented cbVG purified particles, timelapse microscopy 8 - 18hpi, images taken every 1 h. All timelapse images were acquired with a widefield microscope at 20x magnification.

In these experiments, we observed that the signal for the viral reporter gene miRFP670 was low in SG positive cells, to the point where some cells appeared uninfected. This is similar, but more extreme, than our observation via immunofluorescence that SeV NP positive SG positive cells often showed lower signal for SeV NP compared to those that were SG negative cells (**Figure 2A**). We observed similar findings in RSV cbVG-high infection when staining for the RSV F protein (**Figure S4**). We hypothesized that a single cell could gain and lose miRFP670 signal within a 6h window, resulting in SG positive cells that appeared uninfected at the time of imaging. To confirm that SG positive cells during live imaging were infected, we performed timelapse imaging starting at 6 hpi before we begin to see SG positive cells during the infection and tracked SG positive cells every 30 min from 6 to 24 hpi to assess changes in the miRFP670 signal with a higher temporal resolution. SG positive cells showed miRFP670 before forming SG and lost the signal as time went by, demonstrating that SG formation is correlated with a reduction in viral protein expression (**Movie 2 and Figure 7C)**.

### cbVG-mediated interference with viral protein expression is independent on MAVS signaling

The reduction on virus protein levels in SG positive cells led us to hypothesize that the well-established virus interference function of cbVGs is at least in part mediated by the induction of the cellular stress response. Because cbVGs are known to interfere with virus replication through the induction of MAVS signaling and IFN production which consequently leads to a reduction of viral protein levels, we determined if this viral protein reduction observed in SG positive cells was a due to the IFN response and independent on SG formation. To test this, we infected MAVS KO cells with SeV cbVG-high and compared viral protein SeV NP expression to control infected cells. SG positive cells showed similar SeV NP fluorescence in control and MAVS KO cells (**Figure 8A**). These data suggest that the interference in viral protein expression observed in cbVG and SG positive cells is not due to the IFN response, and instead suggest a direct role for the cellular stress response in reducing viral protein expression.

**Figure 8.**
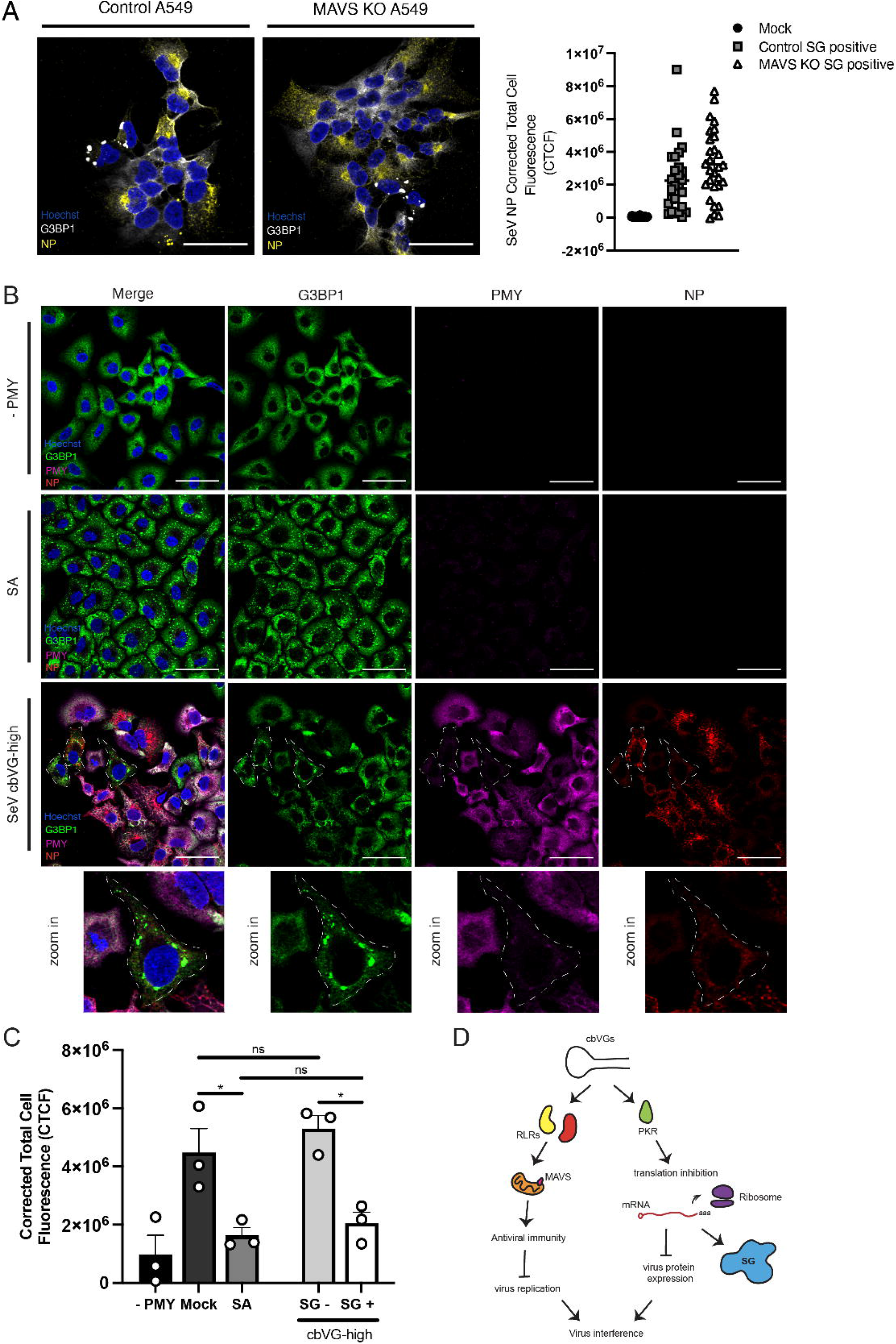
cbVGs induce translation inhibition in SG positive cells. (A) SG (G3BP1 white) and viral protein (SeV NP) detection in control and MAVS KO A549 cells 24 hpi with SeV cbVG-high virus at MOI 1.5 TCID_50_/cell 24 hpi. Corrected total cell fluorescence (CTCF) quantification of SeV viral protein NP in control and MAVS KO A549 SG-positive cells. (B) G3BP1 (green) for SG detection and PMY (magenta) for translation in cells infected with SeV cbVG-high (SeV NP, red) at MOI 3 TCID_50_/cell 24 hpi or treated with sodium arsenite, with and without treatment with PMY for 5 min. (C) Quantification of PMY intensity (Corrected Total Cell Fluorescence) in cells after drug treatment with sodium arsenite or SG positive and SG negative SeV cbVG-high infected cells. Each dot represents the CTCF average of approximately 100 cells counted for each condition. Widefield images were acquired with the Apotome 2.0 at 63x magnification, scale bar = 50 μm. Statistical analysis: one way ANOVA (* p < 0.05)

### cbVGs induce translation arrest in SG positive cells leading to reduced viral protein expression

SG form because of translation inhibition, which could affect virus protein levels in SG positive cells. To determine if translation is inhibited specifically in cbVG-induced SG positive cells, we performed a ribopuromycylation assay to detect active translation at a single cell level using puromycin (PMY) immunostaining. PMY mimics the tyrosine-modified tRNA and upon exposure to cells, is added to nascent peptides. By combining with a translation elongation inhibitor to trap peptides into ribosomes and upon fixing and immunostaining with a PMY-specific antibody, we can detect translation at a single cell level. We performed ribopuromycylation in SeV cbVG-high infected cells and compared PMY staining in SG-positive cells to SG-negative cells. A reduction of PMY signal was observed almost exclusively in SG positive cells during SeV cbVG-high infection (**Figure 8B, lower panel and Figure 8C**). This reduction in signal was comparable to sodium arsenite treated cells (**Figure 8B, middle panel and Figure 8C**). To determine if SG formation is necessary for translation inhibition and reduced viral protein expression, we performed ribopuromycylation in G3BP1/2 dKO and compared PMY staining with control infected cells. As expected, we observe low PMY in G3BP1/2 dKO single cells (**Figure S5**), demonstrating that SG form as a consequence of translation inhibition and are not the drivers of translational arrest. Overall, these data highlight a new function of cbVGs in triggering translation inhibition and SG formation independent of their role in inducing the antiviral response.

## Discussion

cbVGs shape the outcome of SeV and RSV infections (4–8, 34). Their importance is highlighted by their involvement in inducing antiviral immunity, interfering with virus replication, and establishing persistent viral infections (6–8). Here we demonstrate yet another role for cbVGs: to induce translation inhibition by activating PKR signaling and SG formation. PKR activation by cbVGs is independent of the MAVS signaling pathway, highlighting the ability of cbVGs to induce cellular pathways independent of their immunostimulatory activity. Because the content of cbVGs in viral stocks used in experiments is usually not characterized or reported in the literature, our findings that cbVGs play a critical role in the induction of SG help clarify contradicting evidence in the literature related to SG formation and function.

During virus infection, SG can be inhibited by knocking out PKR, inhibiting both translation inhibition and SG formation, or by knocking out nucleating factors like G3BP1, which only inhibits the physical formation of SG and not translation inhibition (11, 35–37). To test the impact of SG formation during cbVG-high infection, we generated a G3BP1 KO model which was previously shown to be required for SG formation during RSV infection (30). To our surprise, knocking out G3BP1 was not sufficient to inhibit the formation of canonical SG containing TIAR (**Figure 5C**). After demonstrating that knocking out G3BP2 was also insufficient to inhibit SG formation, we developed a G3BP1 and G3BP2 dKO cell line, which successfully stopped SG from forming during RSV cbVG-high infection (**Figure 5D**). Our data agrees with a recent report showing that knocking out both G3BP1 and G3BP2 is required for SG inhibition during viral infections (38). We attribute the contradiction regarding the requirement of G3BP1 for SG formation during infection with mononegavirales to differences in the approaches used to validate the absence of SG, and staining for TIAR represents a good alternative SG marker for this purpose.

The immunostimulatory ability of cbVGs together with published data demonstrating how SG are involved in inducing and sustaining the immune antiviral response (13, 14, 31, 39) led us to hypothesize that cbVGs induced SG to aid with activation of the antiviral response. However, using a G3BP1/2 dKO cell line that successfully impeded the formation of SG, we observed no differences in the induction of the antiviral response relative to the control cell line (**Figure 6**). This applied to both the transcriptional and the translational level of IFN and ISG expression (**Figure 6A-C and 6E-F**). A broader transcriptome analysis confirmed that inhibition of SG during infection does not hamper the antiviral immune response (**Figure 6D**). Certain genes were downregulated in the single G3BP1 or G3BP2 KO which indicate SG-independent roles for these proteins during infection. Indeed, G3BP1 is reported to have roles involving the antiviral response (40). We also determined if the effect the cellular stress response had on the global antiviral response was dependent on the translation inhibition that occurred upstream of SG formation. For this, we used the PKR KO system to look at how expression of antiviral genes and proteins were affected. Similar to the G3BP1/2 dKO system, we observed no differences in IFN and ISG expression in both RSV and SeV infection (**Figure 6G-H and S4**), suggesting that the cellular stress response does not impact the overall antiviral response during RSV or SeV infection.

By performing live-cell imaging of G3BP1-GFP expressing cells infected with a reporter SeV, we uncovered interesting facts about the dynamic formation and disassembly of SG. First, we observed that cbVG-induced SG are dynamic and form asynchronously throughout the infection (**Movie 1**). Although we began observing SG positive cells after 8 hpi, the number of SG positive cells increased over time (**Movie 1**). We could also see some cells disassembling SG throughout the infection (**Figure 7A, series 1 and 3**). These data likely explain why not all cbVG-high cells are SG positive when we look at a singular time point 24 hpi (**Figure 1E, 1F**). Although we do not have a working tool in our lab to perform RNA FISH in live cells, we suspect that all cbVG-high cells form SG at some point during the infection, but these are not all captured in snapshot immunofluorescence experiments. The mechanisms governing timing and maintenance of SG formation in a cbVG-positive cell remains unknown. We speculate that there could be a threshold level of cbVG amounts inside the cell that is required to trigger SG formation. Perhaps cell state factors could also be involved in controlling SG formation, such as cell cycle state, circadian rhythm phase, or even metabolic state.

Throughout our studies, we observed a drastic reduction in viral protein level in SG positive cells (**Movie 1 and Figures 2A, S3**). We reasoned that the reduction in virus protein expression could be explained by the well-known function of cbVGs in interfering with the virus life cycle via MAVS signaling and expression of antiviral proteins. In contrast, we saw a reduction of virus protein levels even MAVS KO SG positive cells (**Figure 8A**), suggesting that the interference in protein expression observed in SG positive cbVG-high cells is regulated by activation of the stress response itself. Indeed, we observe drastic translation reduction at a single cell level in SG positive cells infected with cbVG-high virus (**Figure 8B, 8C**). These data implicate translation inhibition and SG formation as a previously undescribed mechanism of cbVG-mediated virus interference. We do not see differences in *RSV G* mRNA in control and PKR KO cells (**Figure 4C**), suggesting PKR activation and translation inhibition does not interfere with virus transcription and instead, interferes with the virus at the translational level due to the translation arrest accompanied by SG formation. We cannot, however, discard a potential additional role for cbVGs in directly interfering with the virus by competing with the virus polymerase, thereby reducing replication, transcription, and translation. However, our group has previously shown this mechanism of interference is minimal (6). Finally, detection of SG in cells several days after infection extends the role of translation inhibition and SG formation to later phases of the infection (**Figure 7B**). It remains unknown if and which other proteins are affected by cbVG-dependent translation inhibition. Although we cannot rule out that a reduction on ISG protein expression occurs specifically in SG positive cells, this potential reduction does not affect the global immune antiviral response during the infection, contradicting reports of a requirement for SG for the initiation of antiviral immunity (14, 15, 17, 38).

The detailed effects of virus protein reduction in RSV and SeV cbVG-high infections remain to be defined, as does the detailed mechanism of how cbVGs induce translation inhibition and SG formation. As SG positive cells are only 10 to 20% of the infected population at a given time, developing tools to isolate and sort SG positive cells from the rest of the population is essential to further understand the role translation inhibition and SG formation play.

Our work highlights how cbVGs, a subset of the virus genome population, are responsible for shaping the cellular stress response during negative-sense RNA virus infection. Although it was previously thought that SG played a role in inducing the global intrinsic innate immune response, our data instead suggest a role for SG in interfering with viral protein translation because of translation inhibition in SeV and RSV infections with high levels of cbVGs. This role extends to later phases of the infection and suggests translation inhibition and SG formation are important both in acute and persistent infections.

## Materials and methods

### Cell lines and viruses

A549 (human type II pneumocytes; ATCC CCL-185) cells were cultured in tissue culture medium (Dubelcco’s modified Eagle’s medium [DMEM; Invitrogen]) supplemented with 10% fetal bovine serum (FBS), gentamicin 50 ng/ml (Thermofisher), L-glutamine 2 mM (Invitrogen) and sodium pyruvate 1 mM (Invitrogen) at 5% CO_2_ 37 °C. The generation of A549 CRISPR KO cell lines has been described previously (41, 42). Plasmids used for CRISPR KO cells lines lentiCRISPR v2 and lentiCRISPR v2-Blast were originally from Feng Zhang (Addgene plasmid # 52961; http://n2t.net/addgene:52961; RRID: Addgene_52961) and Mahan Babu (Addgene plasmid # 83480; http://n2t.net/addgene: 83480; RRID: Addgene_83480), respectively. All KO cell lines were single cell cloned and confirmed KO for each clone was tested through western blot analysis. For generation of A549 G3BP1-GFP, we used the phage UbiC G3BP1-GFP-GFP plasmid originally from Jeffrey Chao (Addgene plasmid # 119950; http://n2t.net/addgene: 119950; RRID: Addgene_119950)(43). G3BP1-GFP expressing cell lines were single cell cloned. Cells were treated with mycoplasma removal agent (MP Biomedical) and tested monthly for mycoplasma contamination using MycoAlert Plus mycoplasma testing kit (Lonza). SeV Cantell strain was grown in 10-day-old, embryonated chicken eggs (Charles River) for 40 h as previously described (44). RSV stocks were grown in Hep2 cells as previously described (18) and harvested by collecting the cells supernatant. SeV and RSV cbVG-high and cbVG-low stocks were produced and characterized as described previously (18).

### Virus infections

For RSV infections, cells were washed once with PBS and then incubated with virus suspended in tissue culture medium supplemented with 2% FBS at 37 °C for 2 h. Cells were then supplemented with additional 2% FBS tissue culture medium. For SeV infections, cells were washed twice with PBS and then incubated with virus suspended in infectious medium (DMEM; Invitrogen) supplemented with 35% bovine serum albumin (BSA; Sigma-Aldrich), penicillin-streptomycin (Invitrogen) and 5% NaHCO_3_ (Sigma-Aldrich) at 37 °C for 1 h. Cells were then supplemented with additional infectious medium. SeV cbVG particles were purified from the allantoic fluid of SeV infected embryonated eggs by density ultracentrifugation on a 5 to 45% sucrose gradient, as described previously (7).

### In Vitro Transcription of cbVGs

The pSL1180 plasmid was cloned to encode the SeV or RSV cbVGs as previously described (9). The cbVG plasmid was linearized, and in vitro transcribed using the MEGAscript T7 kit (Thermofisher). The resulting product was DNAse-treated and purified by LiCl precipitation according to the manufacturer’s protocol. All IVT RNA was quantified by Qubit (Thermofisher) and quality checked through Bioanalyzer (Agilent) to ensure a single band of the correct RNA length was obtained. For transfections, 5 pmol of the IVT cbVG or Low Molecular Weight polyinosine-polycytidylic acid (poly I:C, InvivoGen) were transfected into control A549 or RNAseL-KO A549 cells. At 6 hours post transfection, cells were fixed, permeabilized, and immunostained as described below for Immunofluorescence.

### Recombinant Sendai virus rCantell-miRF670 rescue

To create the pSL1180-rCantell plasmid, the complete viral genome of the SeV Cantell strain and the necessary regulatory elements were inserted into the pSL1180 vector using SpeI and EcoRI restriction enzymes in the following order: T7 promoter, Hh-Rbz, viral genome, Ribozyme, and T7 terminator. A miRF670 gene was then inserted between the NP and P genes in the pSL1180-rCantell plasmid to create the pSL1180-rCantell-miRF670 plasmid. The non-coding region between the NP and P genes was used to separate the NP, GFP, and P genes. Additional nucleotides were inserted downstream of the miRF670 gene to ensure that the entire genome followed the “rule of six”. The NP, P, and L genes of Cantell were cloned into the pTM1 vector to generate the three helper plasmids. All plasmids were validated by sequencing. The recombinant virus rCantell-miRF670 was produced by co-transfecting pSL1180-rCantell-miRF670 and the three helper plasmids. BSR-T7 cells were transfected with a mixture of plasmids containing 4.0 μg pSL1180-rCantell-miRF670, 1.44 μg pTM-NP, 0.77 μg pTM-P, and 0.07 μg pTM-L using Lipofectamine LTX. After 5 h, the medium was replaced with infection medium containing 1 μg/ml TPCK and the cells were incubated at 37 °C. The expression of miRF670 was monitored daily using fluorescence microscopy. At 4 days post-transfection, the cell cultures were harvested, and the supernatants were used to infect 10-day-old specific-pathogen-free embryonated chicken eggs via the allantoic cavity after repeated freeze-thaw cycles. After 40 h of incubation, the allantoic fluid was collected.

### Immunofluorescence

Cells were seeded at 1 x 10^5^ cells/mL confluency in 1.5 glass coverslips (VWR) a day prior infection or drug treatment. The coverslips were transferred to a fresh plate and washed with PBS. Cells were fixed on the coverslips using 4% paraformaldehyde (EMS) for 15 min. Cells were then permeabilized with 0.2% Triton X-100 (Sigma-Aldrich) for 10 min. Primary and secondary antibodies diluted in 3% FBS were added and incubated for 1 h and 45 min, respectively. The nuclei were stained with a 1:10,000 dilution of Hoechst (Invitrogen) in PBS for 5 min prior to mounting. Coverslips were mounted in slides using Prolong Diamond anti-fade mounting media (ThermoFisher) and curated overnight at room temperature. Antibodies used: SeV NP (clone M73/2, a gift from Alan Portner, directly conjugated with DyLight 594 or 647 *N*-hydroxysuccinimide (NHS) ester (ThermoFisher)), RSV NP (Abcam catalog number ab94806), G3BP (Abcam catalog numbers ab181150 and ab56574), G3BP2 (Cell Signaling catalog number 31799), TIAR (Santa Cruz catalog number sc-398372), TIA1 (Abcam catalog number ab40693), Puromycin (Merck, catalog number MABE343).

### RNA FISH combined with immunofluorescence

Cells were seeded at 1 x 10^5^ cells/mL confluency in 1.5 glass coverslips a day prior infection. The coverslips were transferred to a fresh plate and washed with sterile PBS. Cells were fixed in the coverslips using 4% formaldehyde (ThermoFisher) for 10 min and permeabilized with 70% ethanol for 1 h at room temperature. Cells were incubated with primary and secondary antibodies diluted in 1% BSA (Sigma-Aldrich) containing RNAse OUT (ThermoFisher) for 45 and 40 min, respectively. Cells were post fixed with 4% formaldehyde and washed with 2x SSC (ThermoFisher) followed by wash buffer (2x SSC and 10% formamide in water). Cells were hybridized with 2.5 nM RSV specific LGC Biosearch custom probes (See table 1) conjugated to Quasar 570 or Quasar 670. Slides were incubated overnight at 37 °C in a humidified chamber for hybridization. Cells were washed twice with wash buffer for 30 min each and once with 2x SSC for 5 min. Coverslips were mounted using ProLong Diamond Antifade mounting media and curated overnight. Slides were imaged using Zeiss Axio observer widefield microscope.

**Table 1.**
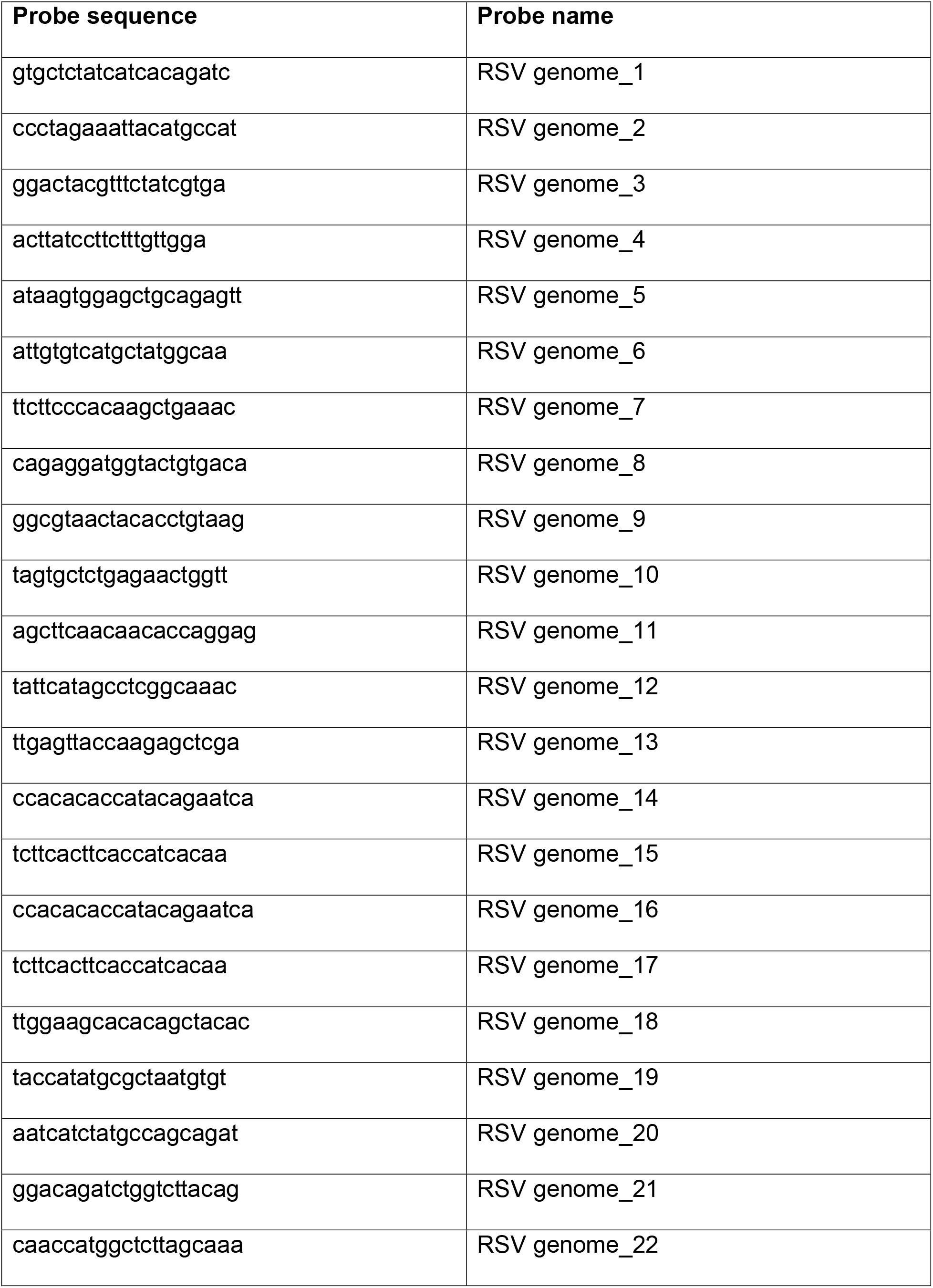
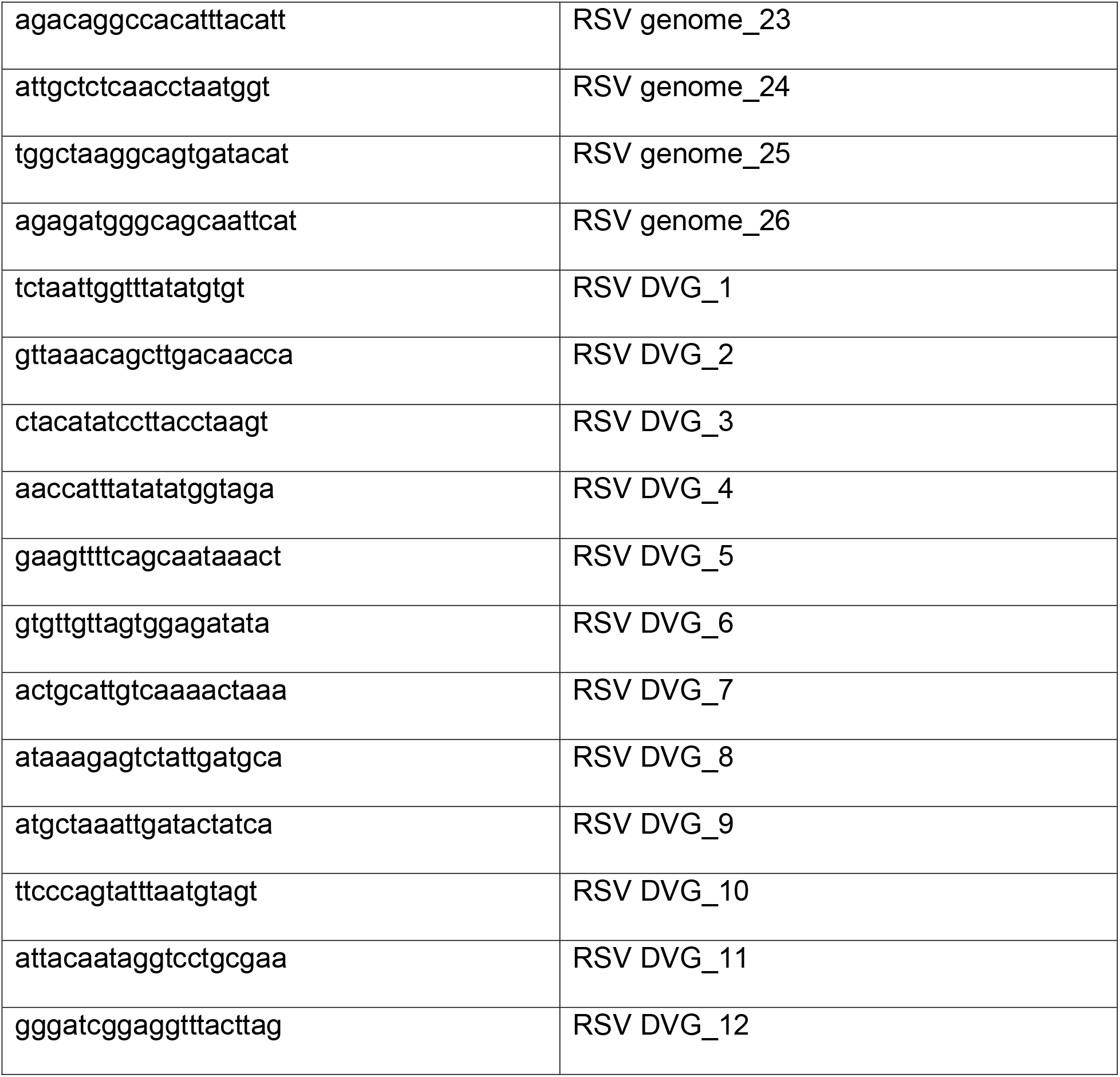
Probe sequences for RSV negative-sense RNA genome probes.

### RNA extraction and PCR/qPCR

RNA was extracted using TriZol reagent (Life Technologies] For qPCR, mRNA was reverse transcribed using high-capacity RNA to cDNA kit (ThermoFisher). qPCR was performed using SYBR green (ThermoFisher) and 5 μM of reverse and forward primers for genes *IL-29* (CGCCTTGGAAGAGTCACTCA and GAAGCCTCAGGTCCCAATTC); *ISG56* (GGATTCTGTACAATACACTAGAAACCA and CTTTTGGTTACTTTTCCCCTATCC); *IRF7* (GATCCAGTCCCAACCAAGG and TCTACTGCCCACCCGTACA) and *RSV G* (AACATACCTGACCCAGAATC and GGTCTTGACTGTTGTAGATTGCA) on an Applied Biosystems QuantStudio 5 machine. Relative mRNA copy numbers were calculated using relative delta CT values and normalized using a housekeeping index with GAPDH and β-actin. For PCR detection of cbVGs, viral RNA was reverse transcribed using a SuperScript III first-strand synthesis (Invitrogen) with Primer GGTGAGGAATCTATACGTTATAC for SeV and primer CTTAGGTAAGGATATGTAGATTCTACC for RSV. PCR was then performed with Platinum Taq DNA polymerase (Invitrogen) with the reverse transcription primers and primer ACCAGACAAGAGTTTAAGAGATATGTATT for SeV and primer CCTCCAAGATTAAAATGATAACTTTAGG for RSV. Bands were analyzed using gel electrophoresis.

### Drug treatments

For sodium arsenite treatment, cells were washed once with PBS and replaced with fresh media. 0.5 mM of sodium arsenite (Sigma-Aldrich) was added to the media and cells were incubated for 1 h at 37 °C. For ISRIB or CHX treatment, infected cells or cells treated with sodium arsenite were treated with 200 nM of ISRIB (Sigma-Aldrich) or 10 μg/mL of CHX for 1 h at 37 °C.

### Ribopuromycylation

Detection of protein translation at a single cell level was adapted from the previously described puromycylation method (45). In brief, cells were seeded at 1 x 10^5^ cells/mL confluency in 1.5 glass coverslips (VWR) a day prior infection or drug treatment. After 24 hpi or 30 min post drug treatment, the media was replaced with PMY labeling medium containing 91 μM of PMY (Sigma-Aldrich) and 45 μM of emetine (Sigma-Aldrich) in tissue culture media and incubated for 5 min at 37 °C. The cells were placed on ice and washed with 1 mL of ice-cold PBS. For PMY removal, the PBS was replaced with extraction buffer containing 0.015% digitonin (Thermo), 50 mM, Tris-HCl pH 8, 5 mM MgCl_2_, 25 mM KCl, Halt™ Protease Inhibitor Cocktail (Thermo), 10 U/mL RNase Out (Thermo) and incubated for 2 min on ice. The extraction buffer was carefully removed and replaced with ice-cold wash buffer containing 50 mM, Tris-HCl pH 8, 5 mM MgCl_2_, 25 mM KCl, Halt™ Protease Inhibitor Cocktail (Thermo), 10 U/mL RNase Out (Thermo). The cells were then fixed with 4% paraformaldehyde (EMS) for 15 min at room temperature. Finally, immunostaining for PMY together with G3BP1 and virus protein NP was performed following the immunofluorescence protocol described above.

### Imaging analysis

SG quantification was performed using Aggrecount automated image analysis as previously described (46). CTCF was analyzed using Fiji software. In brief, cell boundaries were defined with adjusted thresholds using the G3BP1 signal. Then, the CTCF was calculated using the formula: Integrated Density – (Area of selected cell x Mean fluorescence of background).

### Western blots

Cells were seeded at 2 x 10^5^ cells/mL confluency in a day prior infection or drug treatment. Protein was extracted using 1% NP 40 (Thermo) with 2 mM EDTA, 150 mM NaCL (Thermo), 5 mM Tris-HCl, 10% glycerol (Sigma-Aldrich), Halt™ Protease Inhibitor Cocktail (Thermo) and PhosSTOP (Sigma-Aldrich). After samples were incubated on ice for 20 mins and centrifuged for 20 min at 4 °C, supernatant was transferred to new tubes and protein concentration was quantified using the Pierce BCA Protein Assay Kit (Thermo). Protein (10-25 μg) was denatured for 5 min at 95 °C, loaded in a 4%-12% Bis Tris gel (Bio-Rad) and transferred to a PVDF membrane (Millipore Sigma). Membranes were incubated overnight with primary antibodies diluted in 5% BSA in TBS (Fisher) with 0.1 % Tween20 (Sigma-Aldrich). Membranes were incubated with secondary antibodies (anti-mouse or anti-rabbit) conjugated with HRP for 1 h in 5% BSA in TBST. Membranes were developed using Lumi-light Western blot substrate (Roche) to detect HRP and a ChemiDoc (Bio-Rad). Antibodies used for western blot: PKR (Cell Signaling catalog number 12297), p-PKR (Abcam catalog number ab32036), MAVS (Cell Signaling catalog number 3993), G3BP (Abcam catalog numbers ab181150 and ab56574), G3BP2 (Cell Signaling catalog number CS 31799), IFIT1 (Cell Signaling catalog number CS 12082S), IRF7 (Cell Signaling catalog number CS 4920S), RIG-I (Santa Cruz catalog number sc-98911, α-tubulin (Abcam catalog number ab52866).

## Statistics

Statistics were calculated using GraphPad Prism. Version 9

Figure S1. **Characterization of RSV cbVG-high and cbVG-low virus stocks**

Figure S2. **SG positive cells do not show MAVS or RIG-I localization in SG during SeV cbVG-high infection**

Figure S3. **The stress response is not required to induce overall antiviral immunity during SeV cbVG-high infection**

Figure S4. **RSV F protein levels are reduced in SG positive cells during RSV cbVG-high infection**

Figure S5. **SG formation is not necessary for translation inhibition during SeV cbVG-high infection**

Movie 1. **SeV cbVG-dependent SG form asynchronously throughout the infection**

Movie 2. **SG positive cells show decreased virus reporter protein expression**

## Acknowledgements

We would like to acknowledge Dr. Susan Weiss (University of Pennsylvania) for providing the PKR and MAVS KO A549 cells, Nicole Rivera-Espinal for performing the imaging experiments for figure S3 and Emna Achouri for data visualization in Figure 6D. This works was supported by the US National Institutes of Health National Institute of Allergy and Infectious Diseases AI137062 and AI134862 to CLB and T32-007317 to LGA and MH.

## Author contributions

LGA: Conceptualization, methodology, experiments, imaging, analysis, manuscript writing and manuscript editing.

YY: Generation of rCantell-miRF670; manuscript review.

MSH: Generation of IVT DDOs and transfection of DDOs; manuscript review.

CBL: Conceptualization, methodology, manuscript writing and manuscript editing.

---

**Figure 1S. Characterization of RSV cbVG-high and cbVG-low virus stocks**. (A) Agarose gel of cbVG PCR amplicons from A549 cells 24 hpi with RSV cbVG-high virus at MOI 1.5 TCID_50_/cell. (B) Expression of *RSV G* and *IL-29* mRNAs in A549 cells 24 hpi with RSV cbVG-high or cbVG-low virus at MOI 1.5 TCID_50_/cell. Statistical analysis: one way ANOVA (* p < 0.05, ** p < 0.01).

**Figure S2. SG positive cells do not show MAVS or RIG-I localization in SG during SeV cbVG-high infection.** (A) SG (G3BP1, magenta) and MAVS (yellow) staining in A549 cells 24 hpi with RSV cbVG-high virus MOI 1.5 TCID_50_/cell. Zoomed in images of SG positive cells are shown on the right with merge and MAVS and G3BP1 single channel. (B) SG (G3BP1, magenta) and RIG-I (yellow) staining in A549 cells 24 hpi with RSV cbVG-high virus MOI 1.5 TCID_50_/cell. Zoomed in images of SG positive cells are shown on the right with merge and RIG-I and G3BP1 single channel. Widefield images at 40x magnification.

**Figure S3. The stress response is dispensable for overall antiviral immunity during SeV cbVG-high infections** (A) Western blot analysis of phosphorylated IRF-3 in control and PKR KO cells 6 and 18 hpi with SeV cbVG-high at MOI 1.5 TCID_50_/cell. (B) Western blot analysis of IFIT1 in control and PKR KO cells 24 hpi with SeV cbVG-high at MOI 1.5 TCID_50_/cell.

**Figure S4. SG positive cells show reduced RSV F protein expression during RSV cbVG-high infection.** (A) SG (G3BP1, magenta) and viral protein (RSV F, yellow) detection in A549 cells 24 hpi with RSV cbVG-high virus at MOI 1.5 TCID_50_/cell. Zoomed in images of SG positive cells are shown on the right with merge and RSV F single channel. Widefield image was acquired with the Apotome 2.0 at 63x magnification, scale bar = 50 μm.

**Figure S5. SG formation is not necessary for translation inhibition during SeV cbVG-high infection.** (A) G3BP1 (green) for SG detection and PMY (red) for translation in control and G3BP1/2 dKO cells infected with SeV cbVG-high (SeV NP, magenta) at MOI 3 TCID_50_/cell 24 hpi.

**Video 1. SeV cbVG-dependent SG form asynchronously throughout the infection.** G3BP1-GFP expressing A549 cells infected with rCantell-miRF670 reporter virus at MOI 3 TCID_50_/cell with 20 HAU of supplemented cbVG purified particles, timelapse microscopy 6 – 72 hpi, images every 6 h at a 20x magnification.

**Video 2. SG positive cells show decreased virus reporter protein expression**. G3BP1-GFP expressing A549 cells infected with rCantell-miRF670 reporter virus at MOI 3 TCID_50_/cell with 20 HAU of supplemented cbVG purified particles, timelapse microscopy 12 – 24 hpi, images taken every 30 min at a 40x magnification using a widefield microscope.

