## Supplementary figures and images for "Copy-back viral genomes induce a cellular stress response that interferes with viral protein expression without affecting antiviral immunity"

### Fig S1

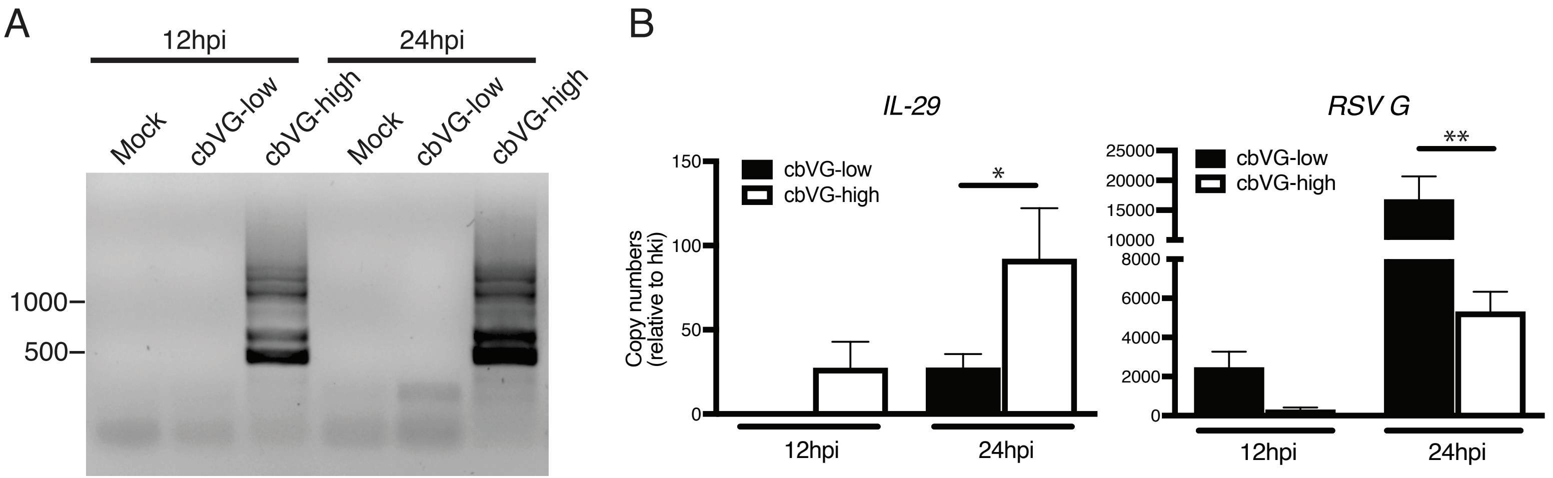

### Fig S2

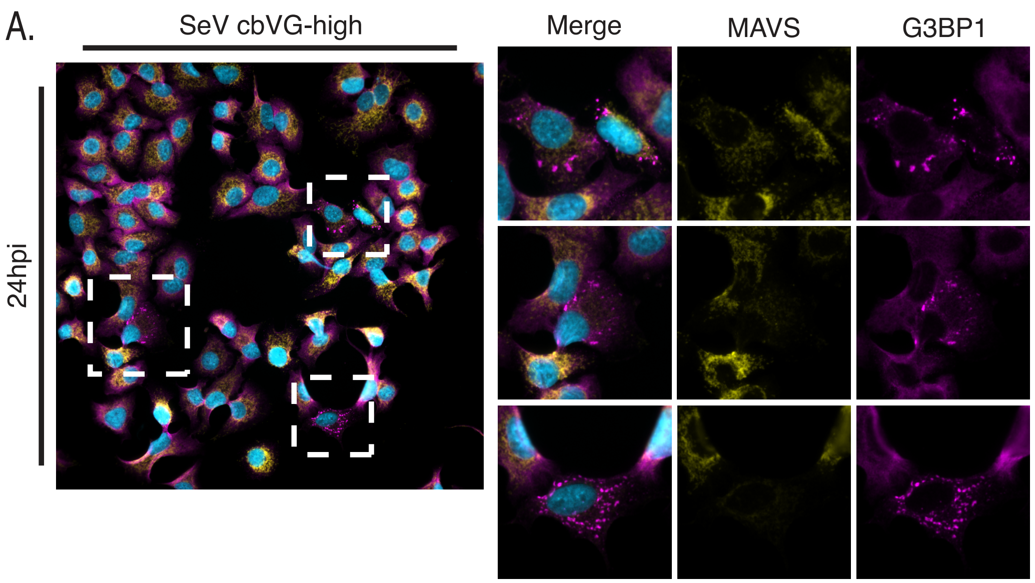

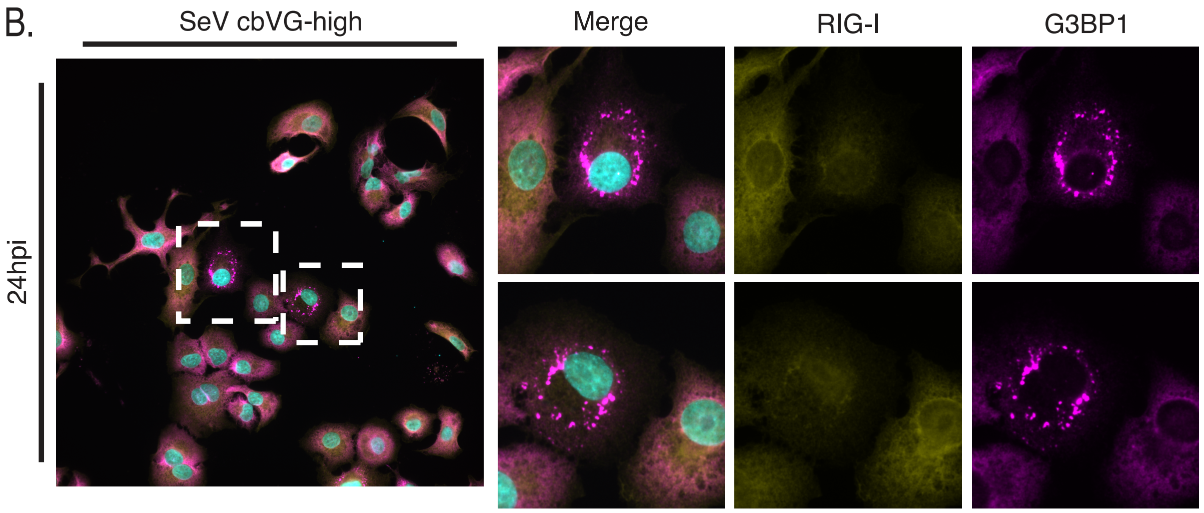

### Fig S3

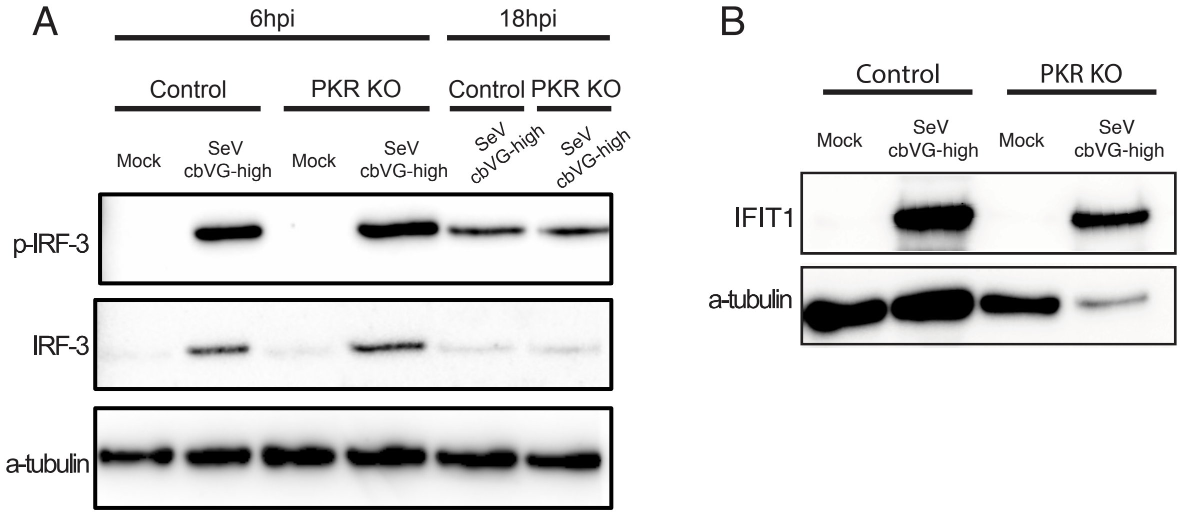

### Fig S4

#### Figure S4

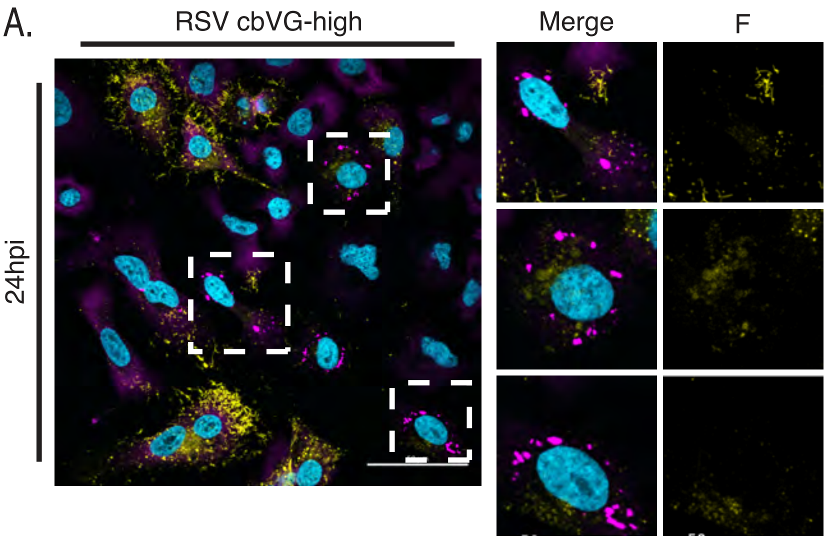
